# Re^2^Pair: Increasing the Scalability of RePair by Decreasing Memory Usage

**DOI:** 10.1101/2024.07.11.603142

**Authors:** Justin Kim, Rahul Varki, Marco Oliva, Christina Boucher

## Abstract

The RePair compression algorithm produces a context-free grammar by iteratively substituting the most frequently occurring pair of consecutive symbols with a new symbol until all consecutive pairs of symbols appear only once in the compressed text. It is widely used in the settings of bioinformatics, machine learning, and information retrieval where random access to the original input text is needed. For example, in pangenomics, RePair is used for random access to a population of genomes. BigRePair improves the scalability of the original RePair algorithm by using Prefix-Free Parsing (PFP) to preprocess the text prior to building the RePair grammar. Despite the efficiency of PFP on repetitive text, there is a scalability issue with the size of the parse which causes a memory bottleneck in BigRePair. In this paper, we design and implement recursive RePair (denoted as Re^2^Pair), which builds the RePair grammar using recursive PFP. Our novel algorithm faces the challenge of constructing the RePair grammar without direct access to the parse of text, relying solely on the dictionary of the text and the parse and dictionary of the parse of the text. We compare Re^2^Pair to BigRePair using SARS-CoV-2 haplotypes and haplotypes from the 1000 Genomes Project. We show that our method Re^2^Pair achieves over a 40% peak memory reduction and a speed up ranging between 12% to 79% compared to BigRePair when compressing the largest input texts in all experiments. Re^2^Pair is made publicly available under the GNU public license here: https://github.com/jkim210/Recursive-RePair

**2012 ACM Subject Classification:** Theory of computation → Formal languages and automata theory

## 1 Introduction

Compressed random access to the text is required for a number of applications including bioinformatics, machine learning, and database access. A *context-free grammar* in Chomsky normal form generates a single string *s* which is referred to as a straight-line program (SLP) for *s*. SLPs prove effective in compactly representing highly repetitive strings with long repeats, as identical substrings can be replaced by a shared non-terminal symbol. Grammar-based compressions use SLPs to represent a string, providing a lossless data compression scheme. Although finding the smallest grammar is generally NP-hard [7, 20], practical polynomial-time compressors like RePair have been proposed.

RePair, pioneered by Larsson and Moffat [12], operates by systematically identifying the most frequently occurring pair of consecutive symbols in a text and replacing it with a novel symbol absent in the original text. This iterative process continues until each pair of consecutive symbols appears only once in the compressed text. While RePair boasts linear time complexity, its notable drawback lies in its substantial space constant, hindering its scalability for large input texts. Although algorithmically simple, RePair achieves high order entropy compression [16], which is why it has been a popular compressor in practice. Since its initial introduction, there have been several different variations and applications of RePair [3, 8, 9, 11]. For example, TreeRePair [13] computes the smallest linear straight-line context-free tree grammar.

BigRePair [10] is another variation that uses Rsync parsing to preprocess the input prior to building the RePair grammar in order to reduce the memory usage of RePair. Notably, the preprocessing that the authors refer to as Rsync parsing is conceptually identical to Prefix-Free Parsing (PFP), a technique which parses the input text *T* in order to build a dictionary (denoted as D_*T*_) and parse (denoted as P_*T*_) of the input. PFP is able to fully represent *T* through D_*T*_ and P_*T*_. PFP was initially devised to efficiently construct the suffix array (SA) and Burrows-Wheeler Transform (BWT) for large repetitive text, but has since found utility across diverse contexts [1, 10, 19]. Its rapid adoption can be attributed to its simplicity and effective scalability for large, repetitive texts. Its efficiency on repetitive text stems from the fact that D_*T*_ exclusively stores non-redundant phrases. This prevents *D*_*T*_ from growing in size when there are repeated phrases in *T*.

A significant drawback of PFP manifests in practice: while the size of D_*T*_ scales admirably for large repetitive text, the scalability of P_*T*_ is less efficient. For instance, Oliva et al.[17] demonstrated that for 2,400 haplotypes of chromosome 19, the parse required over 11 GB of space, whereas the dictionary occupied just over half a GB. Given that P_*T*_ can significantly exceed D_*T*_ in size and often contains repetitions, this naturally suggests that PFP can be effectively applied to P_*T*_ such that P_*T*_ is represented through its dictionary (denoted as DP) and its parse (denoted as PP). This technique of applying PFP to P_*T*_ is referred to as *recursive PFP*. In this paper, we design and implement *recursive RePair* (Re2Pair), a novel algorithm that builds the RePair grammar using recursive PFP.

The algorithmic challenge of this method is building the RePair grammar without direct access to the parse. Instead, we are constricted to constructing the grammar using only the dictionary of the text, and the parse and dictionary of the parse of the text. We note that the main goal of Re2Pair is to significantly reduce the memory footprint of RePair since this is the bottleneck in constructing RePair grammars on large inputs. We first prove (Theorem 4) that Re2Pair uses memory linear in the size of PP, DP, and D_*T*_, which combined is significantly smaller than the size of *T* when *T* is repetitive and not dependent on the size of P_*T*_. Next, we compare our implementation of Re2Pair to BigRePair using SARS-CoV-2 haplotypes and haplotypes from the 1000 Genomes Project [6, 18]. We show that our method Re2Pair achieves over a 40% peak memory reduction compared to BigRePair when compressing the largest input texts in all experiments. As a side-effect, we observe that the RePair component of Re2Pair is faster than that of BigRePair on larger inputs, exhibiting a speed increase ranging from 12% to 79% in our experiments. This speed improvement consistently scales with the input size. Lastly, in our whole genome compression experiment, Re2Pair had the capacity to compress the full 1,200 genome set considered whereas BigRePair could only compress 600 genomes, 50% of Re2Pair capacity, under the same memory limits.

## 2 Preliminaries

### 2.1 Basic definitions

A string *T* is a finite sequence of symbols *T* = *T* [1..*n*] = *T* [1] *T*· · · [*n*] over an alphabet Σ = {*c*_1_, …, *c*_*σ*_} whose symbols can be unambiguously ordered. We refer to the cardinality of the alphabet Σ as the number of symbols in Σ. We denote by *ε* the empty string, and the length of *T* as |*T* |. We denote as *c*^*k*^ the string formed by the character *c* repeated *k* times. We denote by *T* [*i*..*j*] the substring *T* [*i*] · · ·*T* [*j*] of *T* starting in position *i* and ending in position *j*, with *T* [*i*..*j*] = *ε* if *i > j*. For a string *T* and 1≤ *i*≤ *n, T* [1..*i*] is called the *i*-th prefix of *T*, and *T* [*i*..*n*] is called the *i*-th suffix of *T*. We call a prefix *T* [1..*i*] of *T* a *proper prefix* if 1≤ *i < n*. Similarly, we call a suffix *T* [*i*..*n*] of *T* a *proper suffix* if 1 *< i*≤ *n*. Given a set of strings 𝒮, 𝒮 is *prefix-free* if no string in 𝒮 is a prefix of another string in 𝒮. We denote by ≺the lexicographic order: for two strings *T*_2_[1..*m*] and *T*_1_[1..*n*], *T*_2_ ≺*T*_1_ if *T*_2_ is a proper prefix of *T*_1_, or there exists an index 1≤ *i* ≤*n, m* such that *T*_2_[1..*i*− 1] = *T*_1_[1..*i* −1] and *T*_2_[*i*] *< T*_1_[*i*].

### 2.2 Context-free Grammars

A context-free grammar (CFG) is a formal grammar where the rules can be applied to a nonterminal symbol regardless of its context. A CFG is formally defined by a set Σ of terminal symbols, a set *V* of nonterminal symbols, a set ℛ of production rules, and a start symbol 𝒮. A terminal symbol *c* is a symbol that appears in the original text whereas a nonterminal symbol *β* is a new symbol not apart of Σ introduced to the text. A production rule defines how a nonterminal symbol decompresses to a sequence of terminal and nonterminal symbols. Production rules are written in the form of *β*→ *α*, where *α* defines a consecutive sequence of *c* and *β* symbols that appear in the text. The start symbol is defined as the initial nonterminal symbol from which the original text can be reconstructed by applying the production rules. In practice, a CFG is defined by its start symbol and production rules, where the sets of terminal and non-terminal symbols are implicitly defined by these rules.

Chomsky Normal Form is a CFG that requires all production rules adhere to one of the following forms: (1) *β*_*i*_→ *β*_*j*_*β*_*k*_ or (2) *β*_*i*_ →*c*_*i*_ where *β*_*i*_, *β*_*j*_, *β*_*k*_ ∈*V* and *c*_*i*_ ∈ Σ. In other words, a nonterminal symbol should either decompress into (1) two other nonterminal symbols or (2) a single terminal symbol. However, as in other papers [14], we will not explicitly show the production rules for case 2 as those rules are implicitly implied to exist. A straight line program (SLP) is a CFG in Chomsky Normal Form. A SLP is a lossless grammar-based compression scheme representing an input text *T*. With a SLP, random access for any text substring can be achieved with an additive logarithmic time penalty [2, 4]. For brevity, we refer to the SLP that produces an input text *T* simply as a *grammar of T*.

We denote the compressed representation of *T* in the SLP as T.𝒮. Similarly, we denote the set of production rules in the SLP as T. ℛ.

### 2.3 Overview of Prefix-Free Parsing

PFP takes as input a string *T* of length *n*, and two integers greater than 1, which we denote as *w* and *p*. It produces a parse of *T* consisting of overlapping phrases, where each unique phrase is stored in a dictionary. We denote the dictionary as D and the parse as P. We refer to the prefix-free parse of *T* as PFP(*T*). As the name suggests, the parse produced by PFP has the property that none of the suffixes of length greater than *w* of the phrases in D is a prefix of any other. We formalize this property through the following lemma.

#### ▸Lemma 1 ([5]).

*Given a string T and its prefix-free parse PFP* (*T*), *consisting of the dictionary* D *and the parse* P, *the set 𝒮 of distinct proper phrase suffixes of length at least w of the phrases in* D *is a prefix-free set*.

The first step of PFP is to append *w* copies of $ to *T*, where $ is a special symbol lexicographically smaller than any element in Σ with the condition that *T* must not originally contain *w* consecutive copies of $. For the sake of the explanation, we consider the string *T* ^*′*^ = $^*w*^ *T* $^*w*^ 4. Next, we characterize the set of trigger strings E, which define the parse of *T*. Given a parameter *p*, we construct the set of trigger strings by computing the Karp-Rabin hash, *H*_*p*_(*t*), of substrings of length *w* by sliding a window of length *w* over *T* ^*′*^, and letting E be the set of substrings *t* = *T* ^*′*^[*s*..*s* + *w* − 1], where *H*_*p*_(*t*) ≡0 or *t* = $^*w*^. This set E will be used to parse *T* ^*′*^ to find the phrases within the text.

Next, we define the dictionary D and parse P of PFP. Given the string *T* ^*′*^ and a set of trigger strings E, we let D = { *d*_1_, …, *d*_*m*_} be the set of unique phrases in *T* ^*′*^. Each *d*_*i*_ ∈ *D* is a substring of *T* ^*′*^ such that exactly one proper prefix and exactly one proper suffix of *d*_*i*_ is contained within *E* with no other substrings of *d*_*i*_ are allowed to be contained within *E*. The parse *P* contains the ordered occurrences of the phrases that appear in *T* ^*′*^ by pointing to the phrases in the dictionary *D*. We can build *D* and *P* by sliding a window of length *w* across *T* ^*′*^. When a trigger string is encountered, the phrase is added to *D* if not present and the corresponding dictionary phrase pointer is added to *P*. Hence, all phrases start at the beginning of a trigger string and end at the end of the next one. Additionally, the start of the next phrase begins at the start of the last trigger string encountered which means that consecutive phrases in the parse overlap by *w* characters. After scanning *T* ^*′*^, the dictionary is then sorted lexicographically and the pointers in the parse are updated accordingly. We note that *T* can then be reconstructed from D and P alone.

### 2.4 Overview of RePair

RePair creates a context-free grammar from an input text *T* of terminal symbols using the following steps: (1) Find the most frequently occurring pair of consecutive symbols (also known as *bigrams*) within *T*. (2) Replace all occurrences of the most frequently occurring bigram in *T* with a new nonterminal symbol, *β*_*i*_. Repeat these steps until all bigrams in *T* have a frequency of 1; at this point the algorithm halts. The end result of RePair is a grammar (hereon, we denote as *𝒢*) that contains a compressed representation of *T* (hereon, we denote as 𝒮) with no reoccurring bigrams, and a set of rules (hereon, we denote as ℛ) relating every nonterminal symbol to its corresponding pair of symbols. This compressed form is especially desirable because it retains random access to *T* with only a logarithmic time penalty [2, 4]. In particular, *T* can be accessed from ℛ and 𝒮 using the following steps: (1) Read a symbol from 𝒮. If this is a terminal symbol, read the next symbol. Otherwise, replace the nonterminal symbol from 𝒮 with its corresponding pair from ℛ. (2) If there are no more symbols then halt, otherwise repeat from step 1.

The original implementation of RePair has been shown to run in *𝒪* (*n*) expected time and 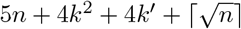 space, where *n* is the number of symbols in *T, k* is the cardinality of Σ, and *k*^*′*^ is the cardinality of the alphabet of the grammar [12]. The most space efficient RePair encoding algorithm to date uses at most *d* log(*d*) + 2*d* bits and runs in *𝒪* (*d*^1.5^) time, where *d* is the number of rules [21]. The most space efficient linear-time RePair encoding algorithm uses (1.5 + *ϵ*)*n* words and runs in *𝒪* (*n/ϵ*) expected time, where *n* is the length of the text and 0 ≤ *ϵ* ≤ 1 [3].

### 2.5 Overview of BigRePair

BigRePair combines PFP with RePair in order to generate a context-free grammar from the input text *T*. Similar to RePair, the BigRePair grammar consists of a set of rules ℛ that relate nonterminal symbols to their corresponding bigrams, and a start sequence 𝒮 that contains nonterminal and terminal symbols with no bigrams occurring more than once. Rather than constructing the grammar directly on the input text *T*, BigRePair begins by running PFP on *T* to generate a dictionary D_*T*_ and parse P_*T*_. The phrases of the dictionary are then concatenated together with a unique separator symbol added between them. We abuse notation for reasons of clarity and denote the resulting string also as D_*T*_.

Next, RePair is ran on P_*T*_ and D_*T*_ to obtain the start sequence and rules for both, which we denote as P_*T*_. 𝒮, P_*T*_. ℛ and D_*T*_. 𝒮, D_*T*_ .ℛ, respectively. We then remove all separator symbols from D_*T*_. 𝒮. Importantly, we note that because unique separator symbols were used to create D_*T*_, there cannot be any rules in D_*T*_. ℛ that contain them as all bigrams including them in *T* must only appear once. Hence, there is nothing to be removed from D_*T*_. ℛ.

Our goal is to create a grammar for *T* (that can be decompressed in accordance with RePair) from the grammars of the dictionary and parse. To obtain *T. 𝒮*, we remind the reader that P_*T*_ defines how the phrases in D_*T*_ need to be placed in relation to one another to create *T*, and similarly, *T. 𝒮* defines the ordering of the rules to generate *T*. It follows then that P_*T*_. 𝒮 defines the start sequence *T*. since P_*T*_. 𝒮 arranges the rules in P_*T*_. ℛ to generate P_*T*_, and P_*T*_ arranges the phrases in D_*T*_ to generate *T*. To obtain *T*., we can combine D_*T*_. 𝒮, D_*T*_. ℛ, and P_*T*_. ℛ. To see this, we first consider naively applying the RePair decompression starting from *T. 𝒮* (which is P_*T*_. 𝒮): we apply the rules of the parse (P_*T*_. ℛ) to obtain the parse P_*T*_, which is a list of the rank’s of the dictionary phrases. At this point, if we could, we would decompress to obtain *T* by simply taking the phrases in D_*T*_ and substituting them into P_*T*_ for their `rank`’s by applying the rules in D_*T*_. ℛ. However, there are two issues: (1) D_*T*_. ℛ does not presently fully encode all the phrases in D_*T*_ in accordance to the RePair decompression scheme. (2) There are no common linkage symbols present between P_*T*_ and D_*T*_, which prevent us from connecting them. To resolve the first issue, for each phrase with `rank` *i* in D_*T*_. 𝒮, we run a variant of RePair where we encode pairs of consecutive symbols into new nonterminals, until the phrases are represented by a single nonterminal, which we denote as *χ*^*i*^. The production rules for these new nonterminals are added to D_*T*_. ℛ. *χ*^*i*^ can then generate the corresponding phrase when it is decompressed in accordance with RePair. Finally, to resolve the second issue, we replace each of the rank’s in P_*T*_. 𝒮 and P_*T*_. ℛ with the corresponding *χ*^*i*^’s.

The motivation behind BigRePair was to use PFP as a preprocessing step to increase the scalability of RePair. However, as mentioned earlier, in practice the size of the parse scales significantly faster than the size of the dictionary. In this next section, we introduce Re2Pair to optimize the size of the parse and further increase scalability.

## 3 Recursive RePair Algorithm

As previously mentioned, BigRePair increases the scalability of RePair for large, repetitive text by using PFP as a preprocessing technique. The scalability of PFP on repetitive text can be broken down by considering the increase in the size of the dictionary and the parse, for increasingly larger input strings. Given an input string *T*, the difference in the size of the dictionary for *T, TT, TTT*, and *TTTT* is negligible in comparison to the size of the parse. Recursive PFP runs PFP on the parse, and then removes the original parse in order to reduce the total size of the output produced by PFP. The challenge reduces to building the original data structure without direct access to the parse.

Here, Re2Pair takes as input *T* and produces a RePair grammar, but builds the grammar using the output of recursive PFP. Our goal is then to build the RePair grammar of *T* (i.e., *T*.ℛ and *T*.𝒮) without having direct access to P_*T*_. We give the details of our solution in this section but preface this discussion with some intuition. We note that if we can obtain P_*T*_. ℛ and P_*T*_. 𝒮 from the recursive PFP components, they can be combined with D_*T*_. ℛ and D_*T*_. 𝒮 to create *T*.R and *T*.S through the method outlined by BigRePair. Further, by accomplishing this, we reduce our goal to creating P_*T*_. ℛ and P_*T*_. 𝒮 using DP. ℛ, DP. 𝒮, PP. ℛ, and PP. 𝒮.

### 3.1 Run Recursive PFP

Re2Pair begins by running PFP on *T*. Algorithm 1 gives the pseudocode for PFP, which has been previously defined in the Preliminaries section. The output is the dictionary D_*T*_ and the parse P_*T*_.

#### Algorithm 1

Prefix-Free Parsing (*w, p, T*)

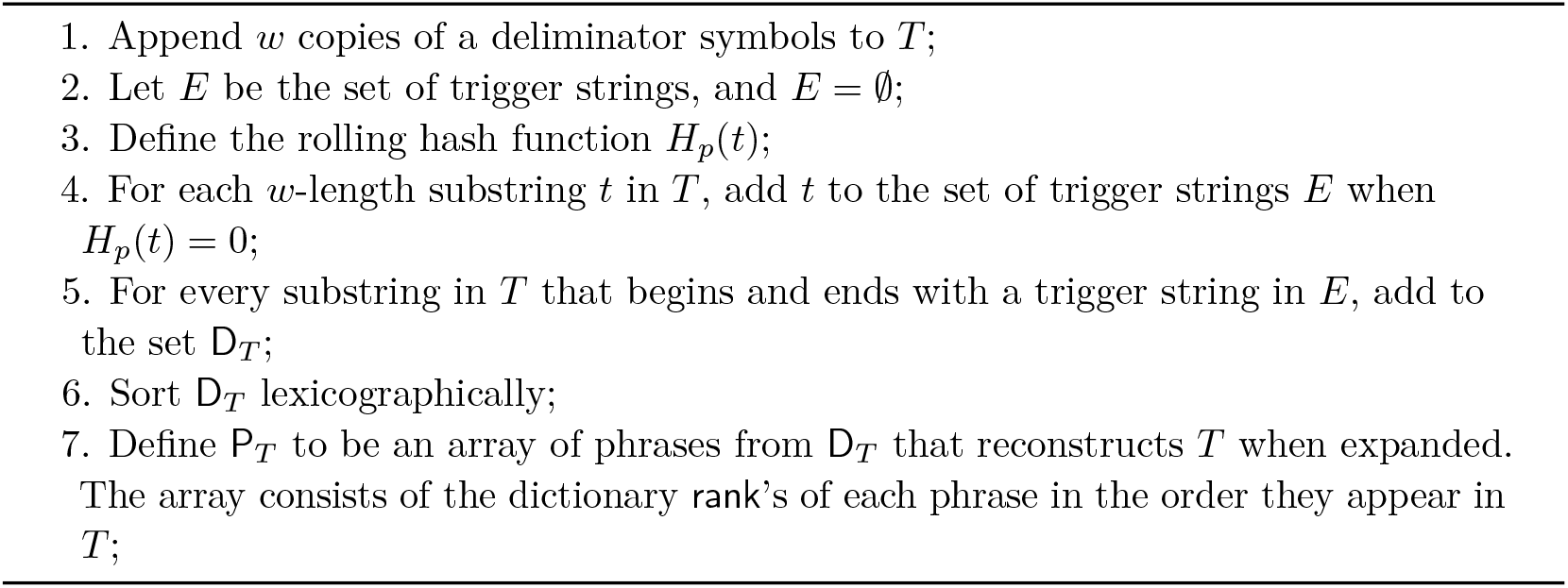

#### ▸Observation 2.

*PFP of T produces P*_*T*_ *and D*_*T*_ *in* 𝒪(|*P*_*T*_ | + |*D*_*T*_ |)*-space*.

Next, recursive PFP is ran, which runs PFP on the parse P_*T*_. The output is the dictionary of the parse, denoted as D_P_, and the parse of the parse, denoted as P_P_. Pseudocode for Recursive PFP is is given in Algorithm 2. The output of recursive PFP is D_*T*_, P_P_ and D_P_.

#### Algorithm 2

Recursive PFP (*w, p*, P_*T*_)

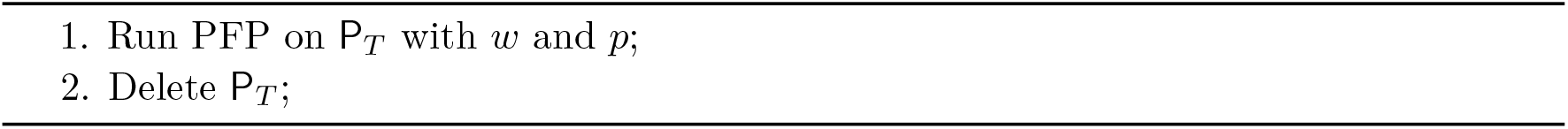

### 3.2 Build Intermediate Grammars Using RePair

In this step, we append a unique separator symbol after each phrase in D_*T*_ and concatenate the phrases into a string. We abuse notation for clarity and refer to the concatenated string also as D_*T*_. We run RePair on this string D_*T*_. Similarly, we do the same procedure for D_P_ to create the concatenated string which we abuse notation and refer to as D_P_, and run RePair on this string D_P_. Lastly, we run RePair on P_P_. After running RePair on each of these strings, we obtain the rules and start sequences for all of them, i.e., P_P_. ℛ and P_P_. 𝒮, D_P_. ℛ and D_P_.𝒮, and D_*T*_. ℛ and D_*T*_. 𝒮. This corresponds to Step 1 of Algorithm 3.

### 3.3 Construct P_*T*_. 𝒮 and P_*T*_. ℛ

We modify D_P_.ℛ, D_P_. 𝒮, P_P_.ℛ, and P_P_.𝒮 in a similar manner as was done in BigRePair in order to construct P_*T*_ .𝒮 and P_*T*_ .ℛ. In particular, we begin by using the unique separator symbols to find the distinct phrases of D_P_ in D_P_.𝒮 Then, we create rules from bigrams within the phrases until only one symbol remains which represents the entire phrase, and add these rules to D_P_.ℛ. We use these symbols to replace the corresponding `rank`’s in P_P_.ℛ and P_P_.𝒮, then concatenate D_P_.ℛ and P_P_.ℛ into P_*T*_ .ℛ and use P_P_.𝒮 as P_*T*_ .𝒮. Next, we demonstrate that this can be accomplished in *𝒪* (|P_P_| + |D_P_| +|D_*T*_|)-space.

The following Observation follows from the fact that the space needed to create the start sequence and rules produced by running BigRePair (PFP + RePair) is bounded by the size of the parse and dictionary of the input.

#### ▸Observation 3.

*Let* P_*T*_ *and* D_*T*_ *be the parse and dictionary obtained from running PFP on T*. P_*T*_. 𝒮, P_*T*_. *R*, D_*T*_ .*𝒮, and* D_*T*_. *ℛ can obtained from running RePair on* P_*T*_ *and* D_*T*_. *Both steps can be done using 𝒪* (|P_*T*_ | + |D_*T*_ |)*-space*.

Next, we make a small observation about the addition of the separators which follows from the fact that the number of separators that need to be added to each dictionary is always less than the cardinality of the respective dictionary.

#### ▸Observation 4.

*Let* D_*T*_ *be the dictionary obtained from running PFP on T, and let* D_P_ *be the dictionary obtained from running recursive PFP. The number of unique separators added to* D_*T*_ *and* D_P_ *does not exceed* |D_*T*_ | *or* |D_P_|, *respectively*.

Using these observations, we can prove bounds on the space needed to construct P_*T*_ .𝒮 and P_*T*_. ℛ. These are given in the following lemmas.

#### ▸Lemma 5.

*Let* D_P_ *and* P_P_ *be obtained by running recursive PFP, and let* D_P_.ℛ, D_P_. *𝒮 and* P_P_. *ℛ be obtained from running RePair on* D_P_ *and* P_P_. *Then* P_*T*_ .*ℛ can be constructed from* D_P_. ℛ, D_P_. *𝒮 and* P_P_.ℛ *in 𝒪* (|P_P_| + |D_P_|)*-space*.

**Proof**. We let D_P_ and P_P_ be obtained by running recursive PFP. Next, we add unique separators in D_P_ to find the distinct phrases in D_P_. It follows that from Observation 4 that the number of unique separators is always less than |D_P_|. We let D_P_.ℛ, D_P_.𝒮 and P_P_.ℛ be obtained from running RePair on D_P_ and P_P_. It follows from Observation 3 that the space needed to generate D_P_.ℛ, D_P_.𝒮 and P_P_.ℛ is at most *𝒪* (|P_P_| + |D_P_|).

Next, we use the new unique separators in D_P_. 𝒮 to find the distinct phrase in D_P_ and for each phrase with rank *i*, we create new rules that generate the phrase from a new nonterminal *χ*^*i*^ (i.e., 2(b) in Algorithm 3). These rules are added to D_P_.ℛ. Hence, there is an addition of at most |D_P_.𝒮| (which is at most |D_P_|) rules to D_P_.ℛ. We remove the separators between phrases in D_P_.𝒮 (i.e., 2(c)), and replace each phrase with `rank` *i* with its corresponding nonterminal *χ*^*i*^ in P_P_.ℛ and P_P_.𝒮. Finally, D_P_.ℛ and P_P_.ℛ are concatenated to create P_*T*_ .ℛ. These last steps do not require additional space, and thus, *𝒪* (|P_P_| + |D_P_|)-space. ◂

#### ▸Lemma 6.

*Let* P_P_ *be obtained from recursive PFP, and* P_P_.*𝒮 be obtained from running RePair on* P_P_. *It follows that* P_*T*_ .*𝒮 can be constructed from* P_P_.*𝒮 in 𝒪* (|P_P_|)*-space*.

**Proof**. We let P_P_ be obtained by running recursive PFP, and P_P_.𝒮 be obtained from running RePair on P_P_, which requires at most *𝒪* (|P_P_|)-space. We obtain P_*T*_ .𝒮 from P_P_.𝒮 by replacing the phrase rank’s in P_P_.𝒮 with the corresponding nonterminals *χ*^*i*^ obtained as described in the proof of Lemma 5. These are one to one substitutions. After this step, P_P_.𝒮 has been converted to P_*T*_ .𝒮. Since the size of P_P_.𝒮 is at most *𝒪* (|P_P_|), it follows that the construction can be accomplished in at most *𝒪* (|P_P_|)-space. ◂

### 3.4 Construct *T. 𝒮* and *T. ℛ*

In the last step of our algorithm (steps 3 and 4 of Algorithm 3), we show that we can construct the BigRePair grammar and rules. This is straightforward since (a) we have previously shown that we can construct P_*T*_. 𝒮 and P_*T*_ .ℛ, (b) we already have D_*T*_. 𝒮 and D_*T*_ .ℛ, and (c) we can now combine (a) and (b) using the BigRePair methodology.

#### ▸Theorem 7.

*We let T be the input text. We let* D_*T*_, D_P_, *and* P_P_ *be obtained from running recursive PFP. We can construct T*.*ℛand T*.*𝒮 from* D_*T*_, D_P_, *and* P_P_ *in 𝒪* (|P_P_|+|D_P_|+|D_*T*_ |)*-space*.

**Proof**. We let D_*T*_, D_P_, and P_P_ be obtained from running recursive PFP. We begin by running RePair on D_*T*_, D_P_, and P_P_ in order to construct D_*T*_ .ℛ, D_*T*_ .𝒮, D_P_.ℛ, D_P_.𝒮, P_P_. ℛ and P_P_. 𝒮. It follows from Gagie et al. [10] that this can be done in *𝒪* (|P_P_| + |D_P_| + |D_*T*_ |)-space. Next, it follows from Lemma 5 that P_*T*_ .ℛ can be constructed from D_P_.ℛ, D_P_.𝒮 and P_P_.ℛ in *𝒪* (|P_P_| + |D_P_|)-space. Further, it follows from Lemma 6 that P_*T*_ .𝒮 can be constructed from P_P_. 𝒮 in *𝒪* (|P_P_|)-space. To construct *T. ℛ* from D_*T*_ .ℛ, D_*T*_ 𝒮. and P_*T*_. ℛ, we first take the unique separators in D_*T*_ .𝒮 to find the distinct phrase in D_*T*_, and for each phrase with `rank` *i*, we create new rules that generates the phrase from a new nonterminal *χ*^*i*^ (i.e., 3(b) in Algorithm 3). These rules are added to D_*T*_ .ℛ. Hence, there is an addition of at most |D_*T*_. 𝒮 | (which is at most |D_*T*_ |) rules to D_*T*_. ℛ. We remove the separators between phrases in D_*T*_ .𝒮 (i.e., 3(c)), and replace each phrase with rank *i* with its corresponding nonterminal *χ*^*i*^ in P_*T*_ .ℛ and P_*T*_ .𝒮. Lastly, we can construct *T*.ℛ by concatenating D_*T*_ .ℛ and P_*T*_ .ℛ, and *T. 𝒮* by letting it be equal to P_*T*_ .𝒮. These last steps do not require any additional space. Therefore, it follows that the complete construction requires at most *𝒪* (|P_P_| + |D_P_| + |D_*T*_ |)-space. ◂

## 4 Experiments

We demonstrate the memory efficiency of Re2Pair by compressing the following datasets: (1) SARS-CoV-2 haplotypes (2) chromosome 1 haplotypes (3) whole genome haplotypes. Specifically, we show that Re2Pair has better memory scalability than BigRePair as the number of sequences in each dataset increases.

### Algorithm 3

Re^2^Pair (D_*T*_, D_P_, P_P_)

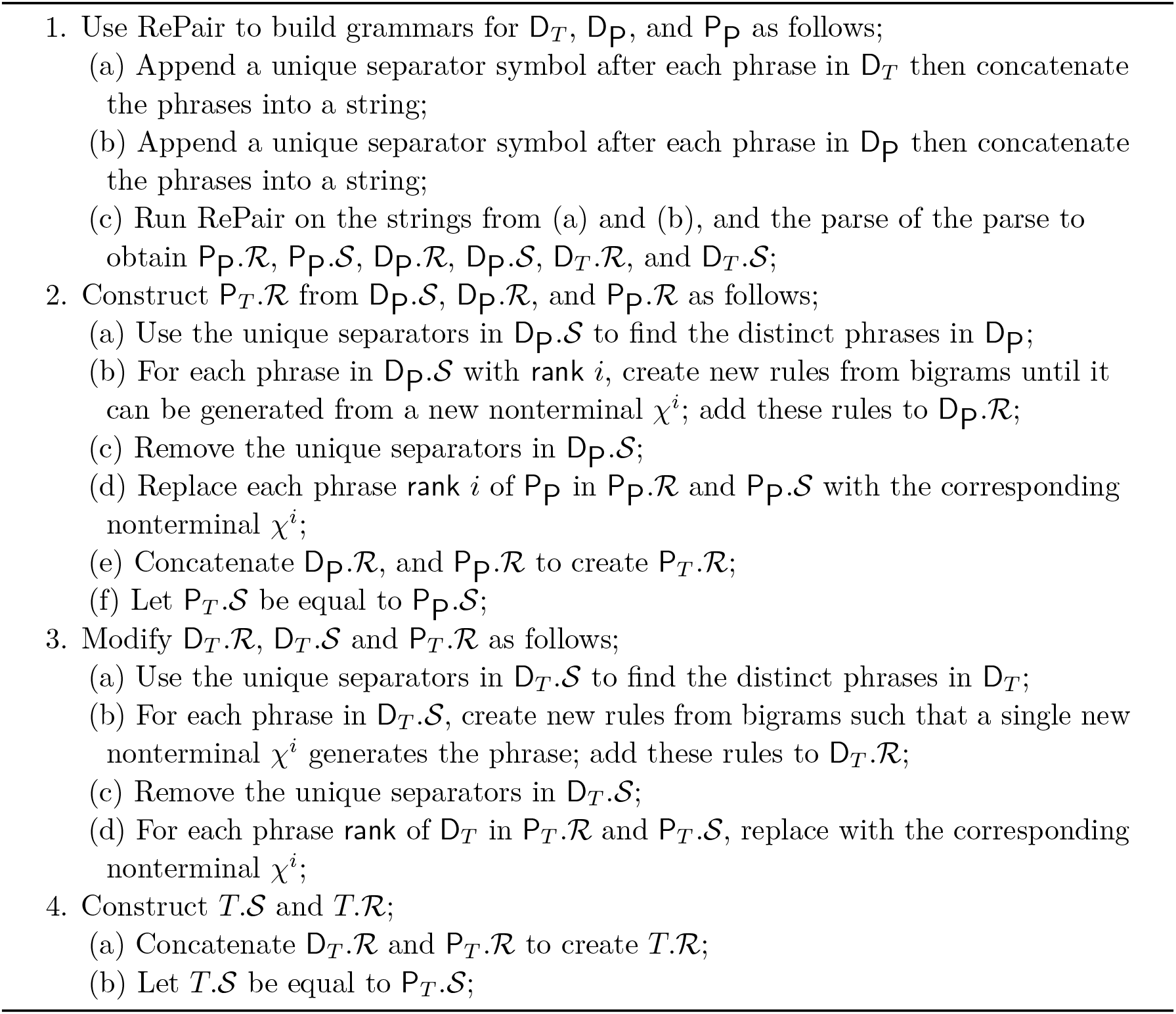

We implemented Re2Pair in ISO/IEC C 9899:2011 and Python. We ran all experiments on a 512 GB server with an AMD EPYC 7702 64-Core Processor running Red Hat Enterprise Linux 8.8. We measured the wall-clock time and peak memory (maximum resident set size) usage of Re2Pair and BigRePair using Snakemake v7.32.4 [15]. As both methods require running PFP on the input data and improving the performance of the PFP step was not our focus, we excluded its time and memory contribution from the reported values for both tools. We note that running PFP on P_*T*_ requires significantly less time and memory compared to running PFP on *T* when *T* is sufficiently large. We ran PFP using 64 threads with its default settings, which set the window size (*w*) to be 10 and the modulus (*p*) to be 100. We ran the RePair portion of both Re2Pair and BigRePair on a single thread since neither tool supports multi-threading for this step currently. We define the compression ratio to be the percentage of the compressed file size (T.𝒮 + T.ℛ) over the uncompressed file size. Experiments that exceeded 512 GB of RAM were omitted from further consideration.

### 4.1 SARS-CoV-2 Genomes

We compared the wall-clock time and peak memory usage of Re2Pair to BigRePair by compressing subsets of SARS-CoV-2 (sars) haplotypes. Specifically, we compressed subsets consisting of 25, 000, 50, 000, 100, 000, 200, 000, and 400, 000 SARS-CoV-2 genomes. Each subset was a superset of the previous one. The smallest subset (`sars.25k`) had an uncompressed size of 0.75GB and the largest subset (`sars.400k`) had an uncompressed size of 12GB. In Table 1, we present the compressed file sizes of these subsets. For the `sars.400k` subset, Re2Pair achieved a compression ratio of approximately 0.29%, slightly higher than the 0.22% ratio achieved by BigRePair.

**Table 1.** The size of the start sequence (S) and rules (R) files generated by BigRePair vs. Re^2^Pair and the corresponding compression ratios for the SARS-CoV-2 subsets compressed. The start sequence and rules file sizes are reported in GB. The uncompressed file size is also reported in GB. Compression ratio is reported as the percentage of the compressed file size over the uncompressed file size.

| Haplotypes | Size | BigRePair |  |  |  | Re<sup>2</sup>Pair |  |  |  |
| --- | --- | --- | --- | --- | --- | --- | --- | --- | --- |
|  |  | S | R | S + R | Ratio | S | R | S + R | Ratio |
| 25,000 | 0.75 | 0.0012 | 0.0024 | 0.0036 | 0.48% | 0.0002 | 0.0045 | 0.0047 | 0.63% |
| 50,000 | 1.61 | 0.0021 | 0.0036 | 0.0057 | 0.35% | 0.0005 | 0.0070 | 0.0075 | 0.47% |
| 100,000 | 3.00 | 0.0034 | 0.0052 | 0.0086 | 0.29% | 0.0010 | 0.0115 | 0.0125 | 0.42% |
| 200,000 | 6.01 | 0.0062 | 0.0087 | 0.0149 | 0.25% | 0.0017 | 0.0199 | 0.0216 | 0.36% |
| 400,000 | 11.88 | 0.0115 | 0.0147 | 0.0262 | 0.22% | 0.0049 | 0.0293 | 0.0342 | 0.29% |

We found that Re2Pair had a peak memory usage that was approximately 43.88% lower than that of BigRePair when compressing the full `sars.400k` haplotype set. Re2Pair required only 1, 551 MiB (1.62 GB) peak memory compared to 2, 764 MiB (2.89 GB) by BigRePair. We see from Figure 1 that Re2Pair begins to use less peak memory than BigRePair at 200, 000 haplotypes. The rate of peak memory growth became slower than BigRePair between compressing the `sars.100k` and `sars.200k` haplotype subsets. A closer examination of the dictionary and parse file sizes revealed that at around 200, 000 haplotypes, the size of the parse started to become larger than the size of the dictionary. This agrees with our experimental results as we only expected Re2Pair to outperform BigRePair in peak memory usage when the size of the parse became larger than the size of the dictionary. If this were not the case, the peak memory usage of both tools would be dictated by running RePair on the dictionary. We note that the wall-clock time was relatively similar between the two methods for all subsets. We saw a modest speed up of 12.53% by Re2Pair compared to BigRePair in compressing the `sars.400k` haplotype set.

**Figure 1.**
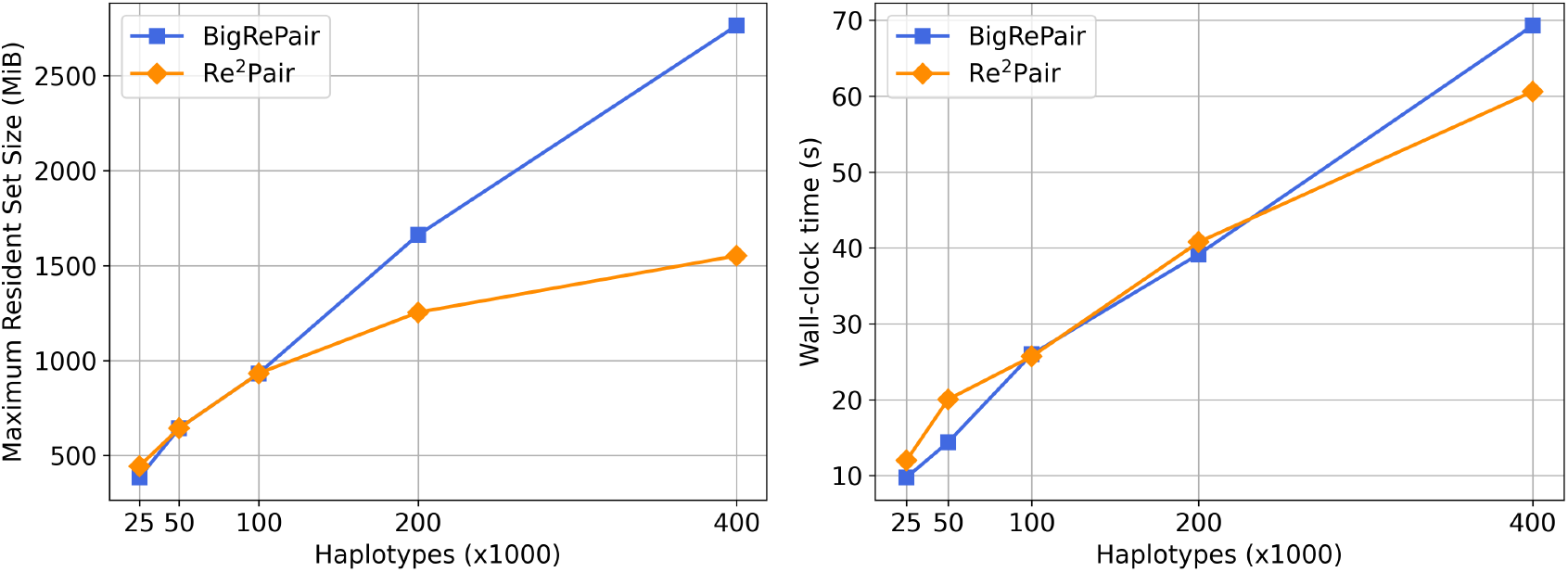
Comparison of the resource usage between BigRePair and Re^2^Pair for compressing the sets of SARS-CoV-2 haplotypes. The left figure shows the peak memory usage (MiB) and the right figure shows the wall-clock time (s) required by each tool.

### 4.2 Chromosome 1 Haplotypes

We next evaluated Re2Pair in comparison to BigRePair by compressing increasingly larger subsets of chromosome 1 (`chr1`) haplotypes from the 1000 Genomes Project. We compressed subsets of chr1 containing 100, 200, 400, 800, 1, 600, and 2, 400 distinct haplotypes. Each subset was a superset of the previous one. The smallest subset (`chr1.100`) had an uncompressed size of 25GB and the largest subset (`chr1.2400`) had an uncompressed size of 600GB. In Table 2, we present the compressed file sizes of these subsets. For the `chr1.2400` subset, Re2Pair achieved a compression ratio of approximately 0.15%, slightly higher than the 0.13% ratio achieved by BigRePair.

**Table 2.** The size of the start sequence (S) and rules (R) files generated by BigRePair vs. Re^2^Pair and the corresponding compression ratios for the chromosome 1 subsets compressed. The start sequence and rules file sizes are reported in GB. The uncompressed file size is also reported in GB. Compression ratio is reported as the percentage of the compressed file size over the uncompressed file size.

| Haplotypes | Size | BigRePair |  |  |  | Re<sup>2</sup>Pair |  |  |  |
| --- | --- | --- | --- | --- | --- | --- | --- | --- | --- |
|  |  | S | R | S + R | Ratio | S | R | S + R | Ratio |
| 100 | 25.09 | 0.01 | 0.25 | 0.26 | 1.04% | 0.005 | 0.28 | 0.28 | 1.12% |
| 200 | 50.17 | 0.02 | 0.27 | 0.30 | 0.60% | 0.01 | 0.31 | 0.32 | 0.64% |
| 400 | 101.32 | 0.04 | 0.30 | 0.34 | 0.34% | 0.02 | 0.36 | 0.37 | 0.37% |
| 800 | 199.89 | 0.07 | 0.36 | 0.43 | 0.22% | 0.03 | 0.46 | 0.50 | 0.25% |
| 1,600 | 401.31 | 0.13 | 0.46 | 0.59 | 0.15% | 0.06 | 0.64 | 0.70 | 0.18% |
| 2,400 | 600.62 | 0.20 | 0.56 | 0.76 | 0.13% | 0.09 | 0.82 | 0.91 | 0.15% |

We see from Figure 2 that Re2Pair compressed all larger subsets of chr1 haplotypes faster and with less memory compared to BigRePair. We found that Re2Pair had a peak memory usage that was approximately 65.22% lower than that of BigRePair when compressing the chr1.2400 haplotype set. Re2Pair required only 50, 815 MiB (53.28 GB) peak memory compared to 146, 143 MiB (153.24 GB) by BigRePair. The peak memory of Re2Pair overall grew 3.33 times slower than that of BigRePair as the number of haplotypes increased. Similarly, we found that Re2Pair was approximately 48.87% faster than BigRePair in compressing the `chr1.2400` haplotype set. Re2Pair required 3, 455 seconds to compress the `chr1.2400` set compared to 6, 719 seconds required by BigRePair.

**Figure 2.**
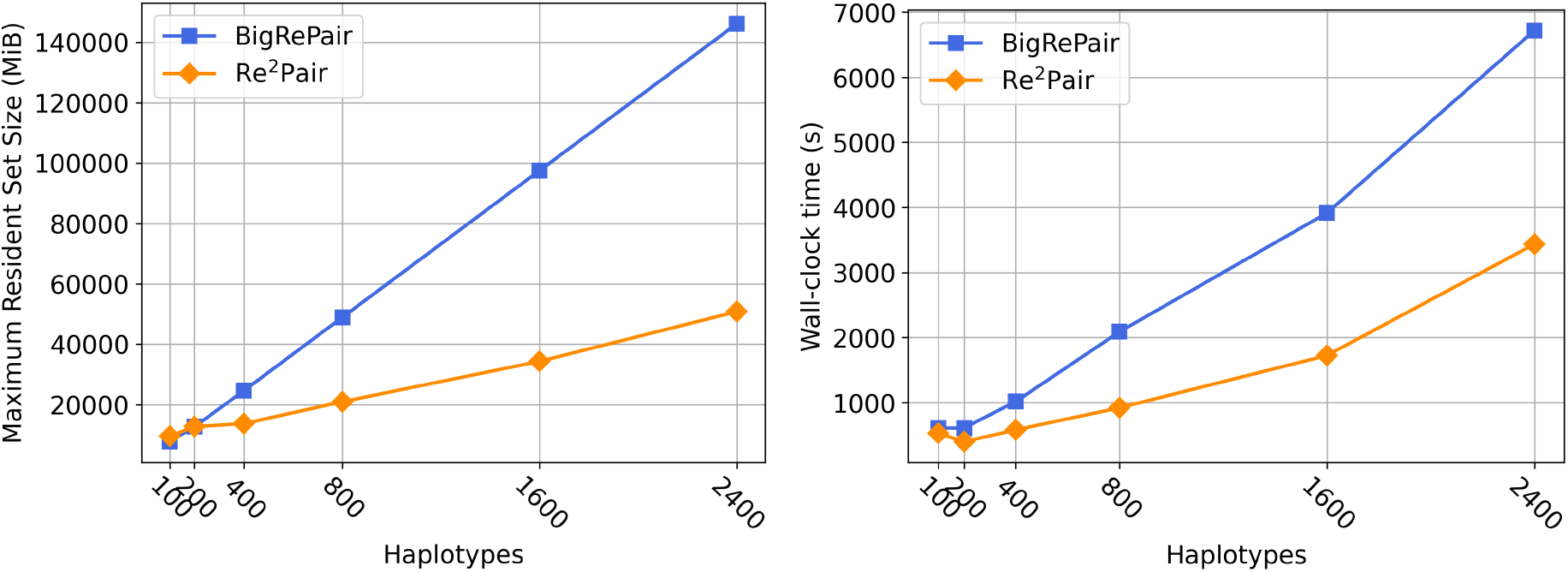
Comparison of the resource usage between BigRePair and Re^2^Pair for compressing the sets of chromosome 1 haplotypes. The left figure shows the peak memory usage (MiB) and the right figure shows the wall-clock time (s) required by each tool.

We see from Table 2 that the size of the start sequence files produced by Re2Pair is less than that of BigRePair for all subsets of haplotypes. Conversely, Re2Pair produced larger rule files than that of BigRePair for all subsets of haplotypes. This makes sense as Re2Pair should produce more compressed start sequences at the cost of more rules compared to BigRePair since the start sequence of Re2Pair is produced by running RePair on the parse of the parse, which is a more compressed version of the parse. We find that the combined size of the start sequence and rules of Re2Pair is slightly worse than that of BigRePair for all inputs. For the largest input (`chr1.2400`), the combined size of the start sequence and rules of Re2Pair was approximately 20% larger than that of BigRePair. We conclude this section by noting that the combined size of the start sequence and rules of Re2Pair for all subsets was less than 1 GB, an insignificant amount of storage by today’s standards.

### 4.3 Whole Genomes from 1000 Genomes Project

Lastly, we compared Re2Pair to BigRePair by compressing increasingly larger subsets of whole genome (`wg`) haplotypes from the 1000 Genomes Project. We define whole genome as containing variants from chr1 to chr22. We attempted to compress the same sets of haplotypes used in the chr1 experiments, namely on the subsets containing 100, 200, 400, 800, 1, 600, and 2, 400 distinct haplotypes. However, the initial PFP step, which is common to both tools, required more than 512 GB to parse the `wg.1600` and `wg.2400` haplotype sets, surpassing the memory limit set. As a result, we had to restrict our interest to subsets containing up to 1, 200 haplotypes, the largest subset that could be evaluated under the memory limit. The smallest subset (`wg.100`) had an uncompressed file size of 295 GB and the largest subset (`wg.1200`) had an uncompressed file size of 3.54 TB. In Table 3, we present the compressed file sizes of these subsets. For the `wg.1200` set, Re2Pair achieved a compression ratio of approximately 0.30%.

**Table 3.** The size of the start sequence (S) and rules (R) files generated by BigRePair vs. Re^2^Pair and the corresponding compression ratios for the whole genome subsets compressed. The start sequence and rules file sizes are reported in GB. The uncompressed file size is also reported in GB. Compression ratio is reported as the percentage of the compressed file size over the uncompressed file size.

| Haplotypes | Size | BigRePair |  |  |  | Re<sup>2</sup>Pair |  |  |  |
| --- | --- | --- | --- | --- | --- | --- | --- | --- | --- |
|  |  | S | R | S + R | Ratio | S | R | S + R | Ratio |
| 100 | 295.16 | 0.18 | 2.79 | 2.97 | 1.00% | 0.06 | 3.11 | 3.17 | 1.07% |
| 200 | 590.71 | 0.30 | 3.00 | 3.30 | 0.56% | 0.11 | 3.44 | 3.55 | 0.60% |
| 400 | 1,180.67 | 0.49 | 3.33 | 3.82 | 0.32% | 0.21 | 4.08 | 4.29 | 0.36% |
| 600 | 1,772.22 | 0.70 | 3.76 | 4.46 | 0.25% | 0.31 | 4.94 | 5.25 | 0.30% |
| 800 | 2,362.98 | - | - | - | - | 0.40 | 6.01 | 6.41 | 0.27% |
| 1200 | 3,540.17 | - | - | - | - | 0.60 | 9.88 | 10.48 | 0.30% |

We see from Figure 3 that Re2Pair outperformed BigRePair in regards to both time and memory for all larger whole genome haplotype subsets. Re2Pair was able to compress the full `wg.1200` haplotype set using 338, 499 MiB (355 GB) peak memory. In comparison, BigRePair was only able to compress up to the `wg.600` haplotype set using 421, 750 MiB (442 GB) peak memory. BigRePair was unable to compress the `wg.800` and `wg.1200` haplotype subsets under the 512 GB memory constraint. Similarly, we found that Re2Pair was 78.95% faster than BigRePair in compressing the `wg.600` haplotype set. Re2Pair required 15, 368 seconds to compress the `wg.600` haplotype set compared to 72, 997 seconds required by BigRePair. We expect that if we allowed BigRePair to exceed the memory threshold and compress the `wg.800` and `wg.1200` haplotype subsets that the time and memory advantage of Re2Pair would have been even larger.

**Figure 3.**
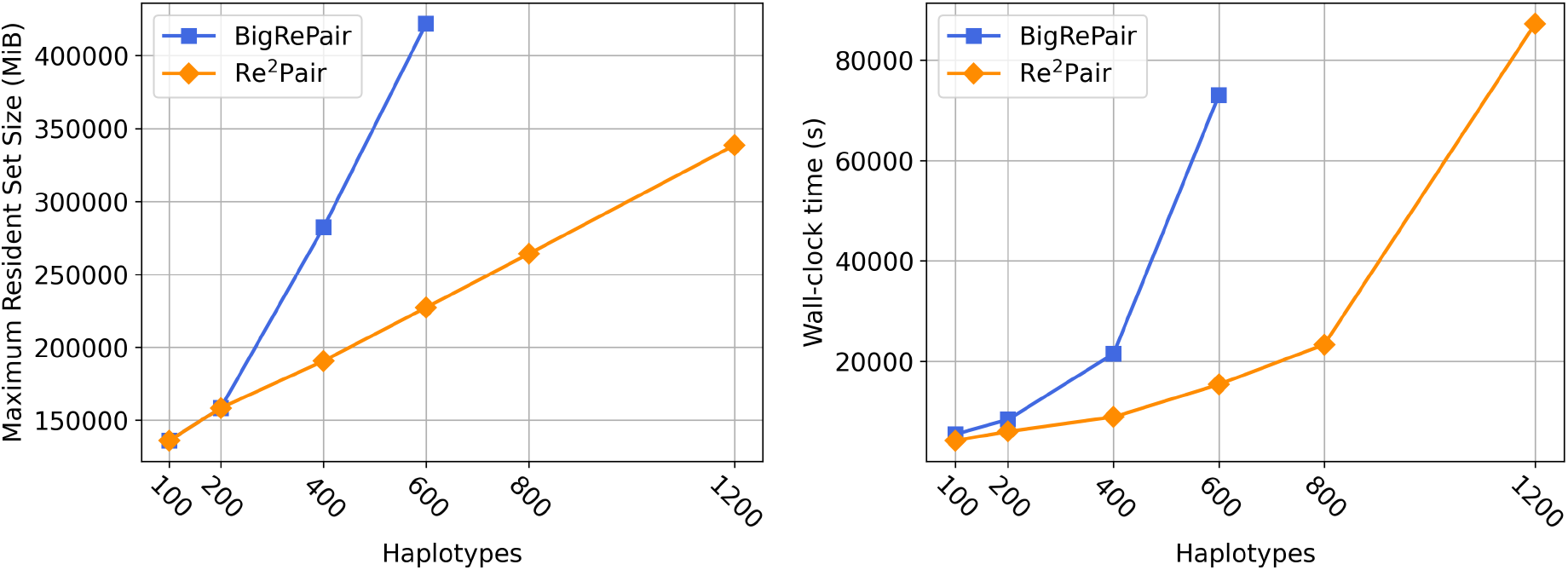
Comparison of the resource usage between BigRePair and Re^2^Pair for compressing the sets of whole genome haplotypes. The left figure shows the peak memory usage (MiB) and the right figure shows the wall-clock time (s) required by each tool.

## 5 Conclusion

When using PFP to index large repetitive datasets, the size of the parse is what causes the memory bottleneck. In this work, we reduce the memory usage of RePair by applying recursive PFP, which runs PFP on the parse of the text. We show that it is possible to build a RePair grammar for an input text using memory proportional to the size of the dictionary of the text and the size of the parse and dictionary of the parse of the text. We prove the correctness of our approach, then run several experiments on real-world datasets that demonstrate the efficacy of our approach in building a RePair grammar on the input text. We observe significant improvements across all experiments when creating a RePair grammar through the recursive PFP components, as opposed to using the PFP components directly. These improvements include over a 40% peak memory reduction and a speed-up ranging between 12% to 79% on the largest input texts.

Finally, we note that recursive PFP has been shown to reduce the memory footprint for two applications of PFP, building the BWT and now building a RePair grammar. We acknowledge that while further PFP recursion on the parse could theoretically reduce the memory usage even more, diminishing returns are likely with each iteration. Applying recursive PFP once has already proven effective in scaling with terabyte size datasets. As datasets grow larger, additional PFP recursion might become necessary. Currently, this technique can be applied on other PFP applications to better optimize their memory usage.

## Digital Object Identifier

10.4230/LIPIcs.ESA.2024.40

## Funding

*Justin Kim*: Completed this project as part of the undergraduate program at the University of Florida.

*Rahul Varki*^1^: [RV was funded by NSF P0183911 SCH: INT: Enabling real time surveillance of antimicrobial resistance given to Dr. Boucher.]

*Marco Oliva*: [MO was funded by NIH R01-AI141810 Developing Computational Methods for Surveillance of Antimicrobial Resistant Agents given to Drs. Boucher and Prosperi.]

*Christina Boucher*: [CB was funded by NIH R01-AI141810 and NSF P0183911.]

## 6 Appendix

### 6.1 PFP Example

We illustrate PFP using a small example. We let *w* = 2 and *T* ^*′*^ = $^2^*T* $^2^ = $$GATTACAT$`GATACAT`$GATTAGATA$$. Now, we assume there exists a Karp-Rabin hash that define the set of trigger strings to be {AC, AG, T$, $$}. It follows that the dictionary D is equal to {$$GATTAC, ACAT$, `AGATA`$$, `T`$`GATAC, T`$`GATTAG` }and the parse P to be [1, 2, 4, 2, 5, 3]

#### 6.2 BigRePair Example

The work in this paper builds upon and is inspired by BigRePair [10]. While the original paper provides valuable insights, we felt that there may have been room for improvement in terms of clarity and organization. A major hurdle in the development of this paper was to clearly understand how the BigRePair algorithm works in practice. We provide here a worked out example of BigRePair for the benefit of the reader to better understand our work in this paper.

We illustrate how BigRePair works with the following example. Suppose we have *T* = ##GATTACAT#AATACAT#AATACATA## with parse P_*T*_ as [1 2 4 2 4 3] (we remove the commas between the rank’s for clarity of the example) and dictionary D_*T*_ as [##GATTAC, ACAT#, ACATA##, T#AATAC]. We apply RePair to the parse P_*T*_ to obtain the start sequence P_*T*_. 𝒮: 1 *α α* 3, and the rules P_*T*_. ℛ : {*α* → 2 4}. Next, we concatenate the phrases of *D*_*T*_ separated by an unique symbol and apply RePair to the final string. In our example, we have the concatenated string: ##`GATTAC`‡`ACAT#`&`ACATA##`$`T#AATAC`, where we color the separator symbols in red for clarity. We apply RePair to this string to obtain the following start sequence and rules.

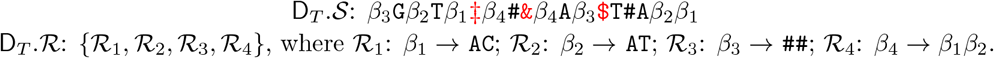

The separator symbols show us the phrase boundaries from D_*T*_ in D_*T*_ .𝒮. Next, we create rules based on bigrams within phrase boundaries until each phrase from D_*T*_ in D_*T*_ .𝒮 is represented by a single nonterminal symbol. Hence, we add the following rules to D_*T*_. ℛ:

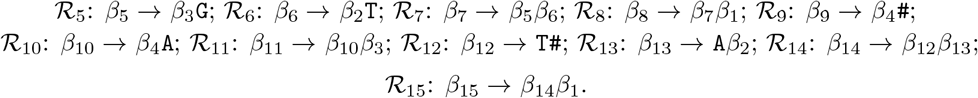

We then add these new rules to D_*T*_. ℛ and apply them to obtain the new D_*T*_ .𝒮.

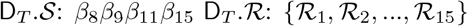

Note that ℛ_5_ − ℛ_8_ decompress *β*_8_ into the first phrase in D_*T*_. 𝒮, ℛ_9_ decompresses *β*_9_ into the second phrase in D_*T*_. 𝒮, ℛ_10_ − ℛ_11_ decompress *β*_11_ into the third phrase in D_*T*_. 𝒮, and ℛ_12_ − ℛ_15_ decompress *β*_15_ into the fourth and final phrase in D_*T*_. 𝒮.

We substitute the rank’s in P_*T*_. 𝒮 and P_*T*_. ℛ with *χ*^*i*^’s. We define nonterminal *χ*^*i*^ for each rank *i* to obtain:

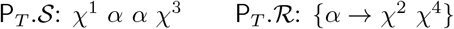

These *χ*^*i*^ are defined to be equal to the symbol which generates a list of nonterminals and terminals that generates a single phrase in *D*_*T*_, essentially taking on the role of a phrase rank. Therefore, we get the following: *χ*^1^ = *β*_8_; *χ*^2^ = *β*_9_; *χ*^3^ = *β*_11_; *χ*^4^ = *β*_15_. The final grammar *T*.*𝒢* consists of the start sequence *T. 𝒮*: *χ*^1^ *α α χ*^3^ and production rules *T. ℛ*: {*α, ℛ*_1_, ℛ_2_, …, ℛ_15_}.

### 6.3 PFP Time and Memory Plots

The following figures show the wall-clock time and peak memory (maximum resident set size) usage of running PFP on the input text *T* and running PFP on the parse of the input text P_*T*_ for the SARS-CoV-2, chromosome 1, and whole genome experiments. We ran PFP on both inputs using 64 threads.

We notice similar trends across all three experiments. The first observation is that the PFP and recursive PFP memory usage is always less than the RePair memory usage on P_*T*_ and D_*T*_. This confirms that the memory bottleneck of BigRePair is in running RePair on P_*T*_ and D_*T*_ rather than running PFP on *T*. The second observation is that the time and memory of running PFP on P_*T*_ is almost always less than running PFP on *T*. The one exception where PFP on P_*T*_ uses more memory than PFP on *T* can be seen in Figure 4. We attribute this observation to the relatively small size of the SARS-CoV-2 dataset, which likely limits the benefits gained from recursive PFP. The last observation is that for the largest subsets of chromosome 1 and whole genome haplotypes compressed, the memory usage of our method Re2Pair, namely running RePair on D_*T*_, P_*P*_, and D_*P*_, uses less memory than running PFP on *T*. With our method, we have experimentally shown that for large datasets we are able to shift the memory bottleneck of the Re2Pair algorithm to be the memory of running PFP rather than the memory of running RePair on the PFP components as is the case with BigRePair.

**Figure 4.**
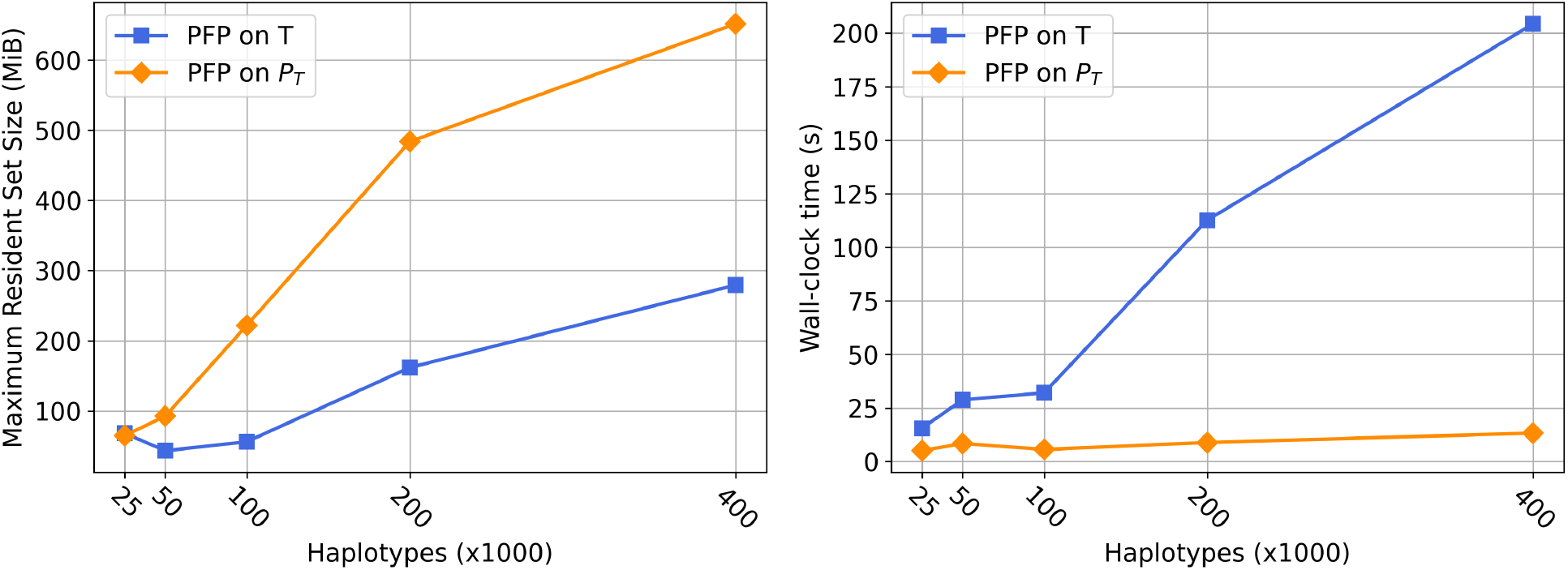
Comparison of the resource usage between running PFP on *T* and PFP on P_*T*_ for the sets of SARS-CoV-2 haplotypes. The left figure shows the peak memory usage (MiB) and the right figure shows the wall-clock time (s) required by each tool.

**Figure 5.**
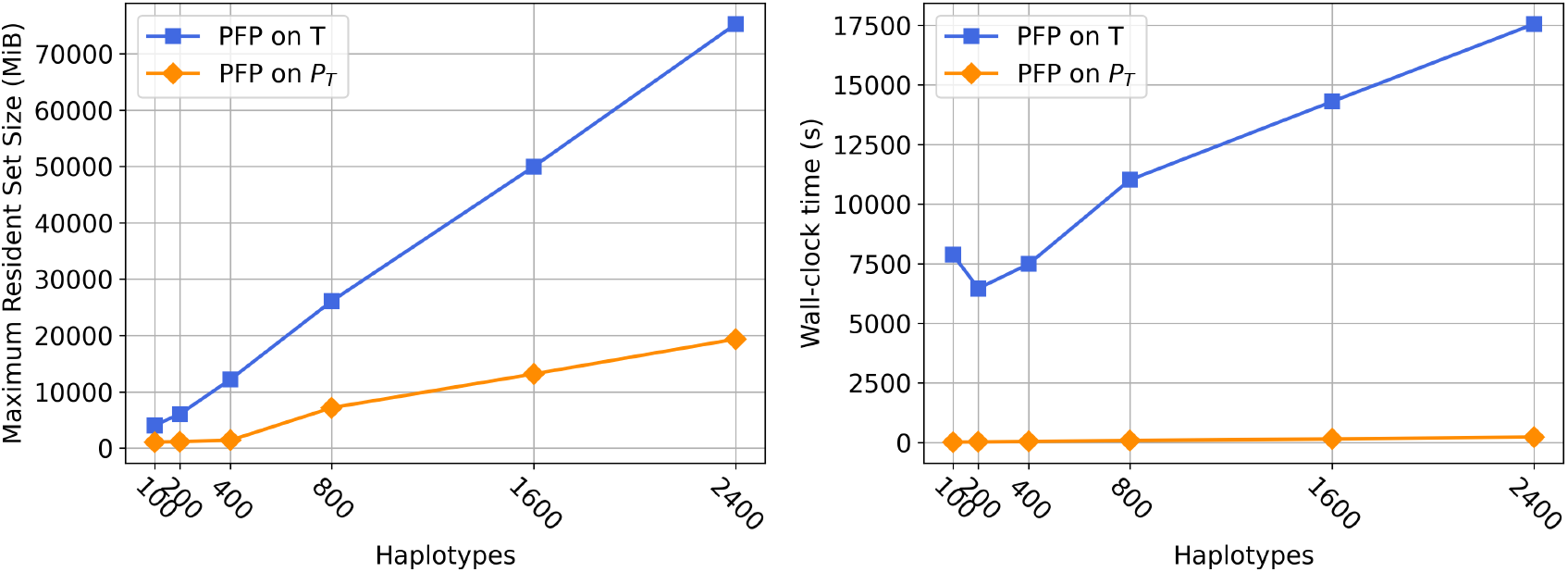
Comparison of the resource usage between running PFP on *T* and PFP on P_*T*_ for the sets of chromosome 1 haplotypes. The left figure shows the peak memory usage (MiB) and the right figure shows the wall-clock time (s) required by each tool.

**Figure 6.**
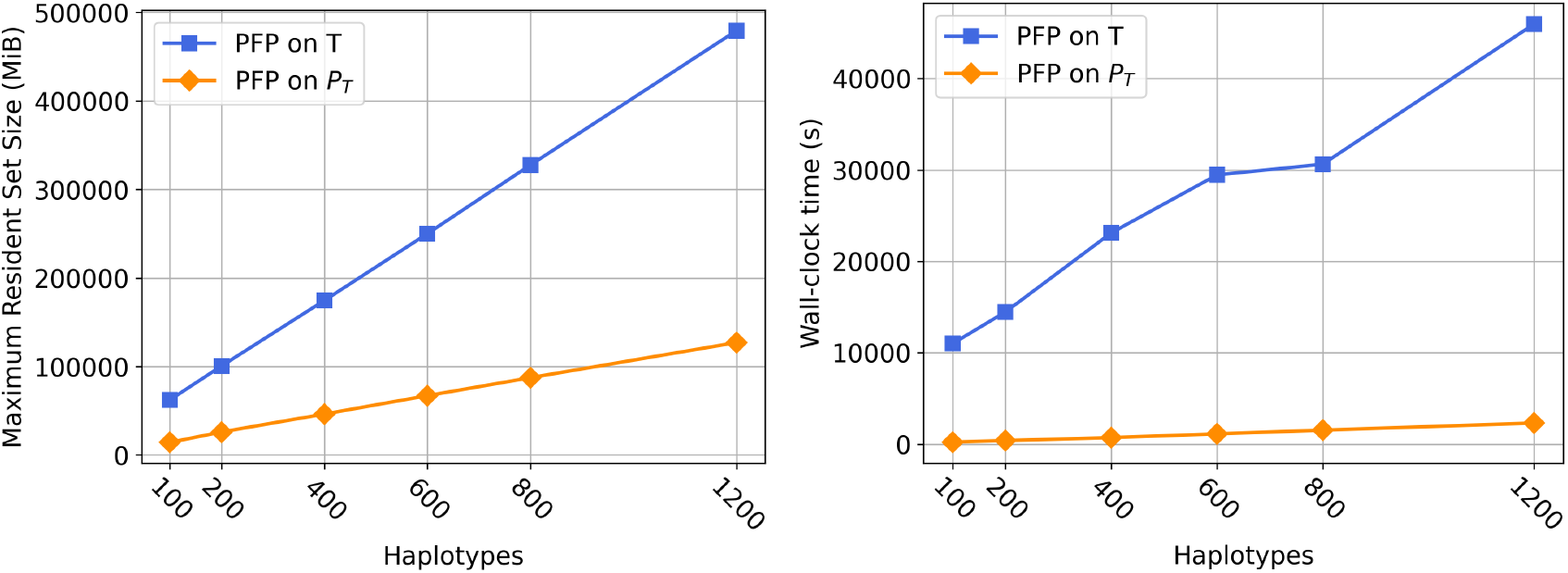
Comparison of the resource usage between running PFP on *T* and PFP on P_*T*_ for the sets of whole genome haplotypes. The left figure shows the peak memory usage (MiB) and the right figure shows the wall-clock time (s) required by each tool.

## Footnotes

4 We note that this definition is equivalent to original definition that considers the string *T*′ = *T*$^*w*^ to be circular.

## Notes

### Competing Interest Statement

The authors have declared no competing interest.

## References

1 Omar Y Ahmed, Massimiliano Rossi, Travis Gagie, Christina Boucher, and Ben Langmead. Spumoni 2: improved classification using a pangenome index of minimizer digests. Genome Biologys, 24(1):122, 2023.

2 Djamal Belazzougui, Patrick Hagge Cording, Simon J Puglisi, and Yasuo Tabei. Access, rank, and select in grammar-compressed strings. In Algorithms-ESA 2015: 23rd Annual European Symposium, Patras, Greece, September 14-16, 2015, Proceedings, pages 142–154. Springer, 2015.

3 Philip Bille, Inge Li Gørtz, and Nicola Prezza. Practical and effective Re-Pair compression. arXiv preprint 1704.08558, 2017.

4 Philip Bille, Gad M Landau, Rajeev Raman, Kunihiko Sadakane, Srinivasa Rao Satti, and Oren Weimann. Random access to grammar-compressed strings and trees. SIAM Journal on Computing, 44(3):513–539, 2015.

5 Christina Boucher, Travis Gagie, Alan Kuhnle, Ben Langmead, Giovanni Manzini, and Taher Mun. Prefix-free parsing for building big BWTs. Algorithms in Molecular Biology, 14(1):13:1–13:15, 2019.

6 Marta Byrska-Bishop, Uday S Evani, Xuefang Zhao, Anna O Basile, Haley J Abel, Allison A Regier, André Corvelo Wayne E Clarke, Rajeeva Musunuri, Kshithija Nagulapalli, et al. High-coverage whole-genome sequencing of the expanded 1000 Genomes Project cohort including 602 trios. Cell, 185(18):3426–3440, 2022.

7 Moses Charikar, Eric Lehman, Ding Liu, Rina Panigrahy, Manoj Prabhakaran, Amit Sahai, and Abhi Shelat. The smallest grammar problem. IEEE Transactions on Information Theory, 51(7):2554–2576, 2005.

8 Francisco Claude, Antonio Farina, Miguel A Martínez-Prieto, and Gonzalo Navarro. Compressed q-gram indexing for highly repetitive biological sequences. In 2010 IEEE International Conference on BioInformatics and BioEngineering, pages 86–91. IEEE, 2010.

9 Francisco Claude and Gonzalo Navarro. Fast and compact web graph representations. ACM Transactions on the Web (TWEB), 4(4):1–31, 2010.

10 Travis Gagie, Tomohiro I, Giovanni Manzini, Gonzalo Navarro, Hiroshi Sakamoto, and Yoshimasa Takabatake. Rpair: Rescaling RePair with Rsync. In International Symposium on String Processing and Information Retrieval, pages 35–44. Springer, 2019.

11 Rodrigo González and Gonzalo Navarro. Compressed text indexes with fast locate. In Annual Symposium on Combinatorial Pattern Matching, pages 216–227. Springer, 2007.

12 N Jesper Larsson and Alistair Moffat. Off-line dictionary-based compression. Proceedings of the IEEE, 88(11):1722–1732, 2000.

13 Markus Lohrey, Sebastian Maneth, and Roy Mennicke. XML tree structure compression using RePair. Information Systems, 38(8):1150–1167, 2013.

14 Takuya Mieno, Shunsuke Inenaga, and Takashi Horiyama. RePair grammars are the smallest grammars for fibonacci words. In 33rd Annual Symposium on Combinatorial Pattern Matching, CPM 2022, Leibniz International Proceedings in Informatics, LIPIcs. Schloss Dagstuhl-Leibniz-Zentrum fur Informatik GmbH, Dagstuhl Publishing, June 2022. doi:10.4230/LIPIcs.CPM.2022.26.

15 Felix Mölder, Kim Philipp Jablonski, Brice Letcher, Michael B Hall, Christopher H Tomkins-Tinch, Vanessa Sochat, Jan Forster, Soohyun Lee, Sven O Twardziok, Alexander Kanitz, et al. Sustainable data analysis with Snakemake. F1000Research, 10, 2021.

16 Gonzalo Navarro and Luís Manuel Silveira Russo. Re-Pair achieves high-order entropy. In DCC, page 537, 2008.

17 Marco Oliva, Travis Gagie, and Christina Boucher. Recursive Prefix-Free Parsing for Building Big BWTs. In 2023 Data Compression Conference (DCC), pages 62–70. IEEE, 2023.

18 Arang Rhie, Sergey Nurk, Monika Cechova, Savannah J Hoyt, Dylan J Taylor, Nicolas Altemose, Paul W Hook, Sergey Koren, Mikko Rautiainen, Ivan A Alexandrov, et al. The complete sequence of a human y chromosome. Nature, 621(7978):344–354, 2023.

19 Massimiliano Rossi, Marco Oliva, Ben Langmead, Travis Gagie, and Christina Boucher. Moni: A pangenomic index for finding maximal exact matches. Journal of Computational Biology, 2022.

20 James A Storer and Thomas G Szymanski. Data compression via textual substitution. Journal of the ACM (JACM), 29(4):928–951, 1982.

21 Yasuo Tabei, Yoshimasa Takabatake, and Hiroshi Sakamoto. A succinct grammar compression. In Annual Symposium on Combinatorial Pattern Matching, pages 235–246. Springer, 2013.

